# LadybirdBase: a comprehensive biology, ecology and omics resource for ladybird beetles (Coccinellidae)

**DOI:** 10.1101/2025.06.15.659824

**Authors:** Yi-Fei Sun, Kun-Yu Yang, Hao Li, Yuan-Sen Liang, Li-Qun Cai, Jia-Yi Xie, You-Wen Zhang, Jia-Yong Liang, Qian Mou, Ying-Min Wang, Dan Chen, Meng-Xue Qi, Luis Carlos Ramos Aguila, Hao-Sen Li, Hong Pang

**Affiliations:** State Key Laboratory of Biocontrol, School of Ecology, Sun Yat-sen University, Shenzhen 518000, China

**Keywords:** Database, Coccinellidae, genome, microbiome, biology, ecology

## Abstract

Ladybird beetles (Coleoptera: Coccinellidae) represent a highly diverse insect group comprising over 6000 species. Recently, an increasing amount of omics data on ladybirds has accumulated. To better understand their roles and adaptations in biological control, we developed LadybirdBase (http://www.ladybirdbase.com/), a comprehensive database that compiles both published and self-collected data on ladybird beetles. It contains five main modules that encompass: (1) the Biology Module summarizing information on over 6000 species, including their classification, morphology, diet, distribution, and omics data; (2) the Ecology Module offering diet and distribution records; (3) the Genomics Module integrating published and unpublished resources such as genomes, transcriptomes, mitogenomes, orthogroups, and variance; (4) the Microbiomics Module presenting 16S rRNA, ITS, and metagenome data; and (5) the Lab Test Module covering diet ranges, life histories under different treatments, and stress tolerance. Additionally, LadybirdBase offers analysis tools for species detection using morphology or barcodes, BLAST searches against the genome resources, differential gene expression analysis, and PCR primer search. LadybirdBase can greatly simplify species identification, enhance the efficiency of omics studies, and provide a theoretical foundation for the use of ladybirds in biological control.

## 1. Introduction

Ladybird beetles (Coleoptera: Coccinellidae) comprise a diverse group of insects with over 6,000 species distributed across all biogeographic regions (1,2). They exhibit a wide range of feeding behaviors—such as coccidophagy, aphidophagy, fungivory, and phytophagy—making some species valuable biological control agents, like *Cryptolaemus montrouzieri, Coccinella septempunctata, Harmonia axyridis*, and *Propylea japonica* (3). However, challenges have emerged in deploying ladybirds as biocontrol agents. One major issue is the lack of detailed knowledge on predator-prey relationships and geographic distributions, which can lead to ineffective releases or even new invasions, as in the case of *H. axyridis*, now considered invasive in many regions (4,5). Another concern is the potential non-target effects of introduced ladybirds, which have been observed or predicted both in laboratory and field studies (6). Additionally, incompatibility between ladybird populations and insecticides can impair key traits such as mobility, reproductive capacity, and predatory efficiency, limiting the combined use of chemical and biological control (7). Moreover, traditional rearing systems for ladybird predators are time-consuming and costly, highlighting the need for developing artificial diets and evaluating their performance (8).

The recent surge in high-throughput sequencing technologies has rapidly expanded genomic and microbiomic resources for insects, enabling studies on their performance, evolutionary dynamics, and trait development (9). Genomics researches provide theoretical basis and the candidate targets for understanding the responses in facing the stresses and their adaption, which can further use in practical production and upgrading species, like the assessment before releases, developing insecticide-resistance strain, and exploiting artificial diet (10). Microbiome researches have further revealed critical roles in host digestion, detoxification, and overall fitness (11,12). Several studies on ladybirds have leveraged these omics resources to explore systematics, phylogeny, gene functions, and adaptive evolution. For example, research on *C. montrouzieri* demonstrated its rapid genomic changes associated with introduction and adaptation to alternative prey (13,14). Microbiome also showed flexibility in response to different conditions (15-18). Despite these advances, researchers face challenges such as fragmented data, limited accessibility, and inconsistent analysis methods, which hinder broader application and integration.

Several ladybird group-specific databases have been development, including Ladybirds of Australia (https://www.ento.csiro.au/biology/ladybirds/ladybirds.htm), Coccinellidae de Chile (https://www.coccinellidae.cl/), and The Lost Ladybug Project (http://www.lostladybug.org). These databases mainly present the data of morphological taxonomy and distribution. Although there are several insect-focused omic databases, such as InsectBase (19), Hymenoptera Genome Database (20), SilkDB (21), no ladybird-specific omic database exist. For example, InsectBase includes only four ladybird species and lacks experimental designs and microbiome data relevant to ladybirds (data accessed March 4, 2025). Therefore, there is a clear need for a comprehensive database dedicated to ladybird beetles that integrates species information, omics data, and relevant studies.

In this study, we present LadybirdBase, a specialized database for ladybird beetles that compiles published references and unpublished data from our lab. LadybirdBase includes modules covering biology and ecology, genomics, microbiomics, and laboratory test data (Figure 1). Additionally, it features tools for species identification, gene BLAST searches, differential gene expression analysis and PCR primer search (Figure 1). By consolidating these resources, LadybirdBase aims to provide a comprehensive, user-friendly data platform that offers accurate information about Coccinellidae, supporting future research and the use of ladybirds in biological control.

**Figure 1.**
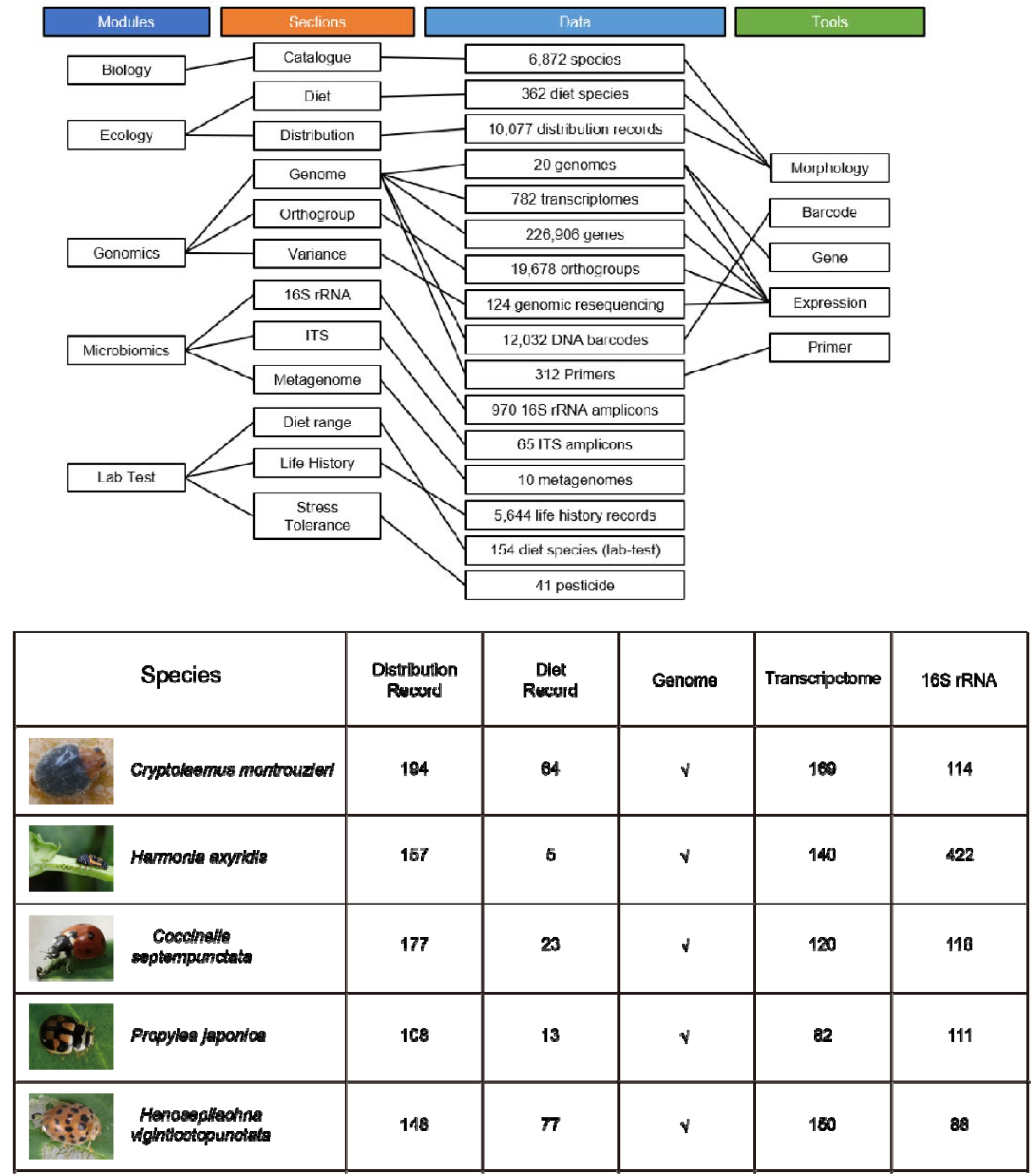
The construction workflow of LadybirdBase.

## 2. Materials and Methods

### 2.1. Module data collection and analysis

For the Biology Module, species information—including taxonomy, key morphological traits, distribution, and diet—was primarily collected from published literature and the Wiki Species website (https://species.wikimedia.org/wiki/Main_Page), with manual verification. The classify of subfamilies and tribes followed the result of Che et al. (22), and genuses followed Tomaszewska et al. (23). Some groups with disputes are specifically pointed out in the ‘introduction’ part.

For the Ecology Module, the information of diet and distribution were collected and displayed in ‘Diet’ and ‘Distribution’ sections. Based on the records finding in the field, the preys are divided into 10 types, including coccids, aphids, whiteflies, mites, plants, thrips, psyllids, ants, planthoppers, and pollen. The distribution of each species, including their countries and regions were accessible.

For the Genomics Module, genomics data were sourced from public databases and partially from unpublished datasets. We included ten assembled genomes of their sequences and the results of gene prediction from our previous comparative genomic study (24) and gathered 685 transcriptome datasets from 17 species, either from the NCBI SRA database or our own unpublished data. All transcriptome data and the partition of orthogroups were analyzed as described in Huang et al. (24). Gene families reported to act in the adaption of dietary, like chemosensory-related, digestion-related, detoxification-related, and immunity-related, were classified in Huang et al. (24). Reported gene functions were compiled from literature and sequences obtained from NCBI. BLAST searches (e-value <1e-5) were performed against the assembled genomes to annotate homologous genes. The detection of potential HGT events in the 10 genomes were based on the HGT index (*h*), which have been used in several studies (25-27). We further excluded contaminations by phylogenies and gene structures. Detail method for HGT detection can be found in Additional File 1.

For the Variance Section of the Genomics Module, genomic resequencing data for *Cryptolaemus montrouzieri* were derived from unpublished datasets. A total of 56 laboratory-reared and 68 field-collected *C. montrouzieri* individuals were subjected to genomic resequencing (approximately 10× coverage on the Illumina NovaSeq 6000). Clean reads were mapped to the reference genome (24) using BWA v0.7.17 (28), and BAM files were processed with SAMtools v1.13 (29) and Picard Tools v2.26.2 (https://broadinstitute.github.io/picard/) for duplicate marking. Variants were called and filtered using GATK v4 (30), applying hard filtering to remove false positives. SNPs were retained if minor allele frequency was ≥0.05, missing rate ≤20%, and maximum allele count was two, ensuring heterozygosity conformed to Hardy–Weinberg expectations. Functional annotations were performed using SnpEff (31).

For the Microbiomics Module, we collected 16S rRNA, ITS amplicon, and metagenomes sequencing datasets from NCBI SRA and unpublished sources. 16S rRNA reads were all analysis using an in-house script based. Quality-filter and process used QIIME2 (32). Divisive Amplicon Denoising Algorithm (DADA2) was employed for denoising and chimera removal (33). Taxonomic classification of ASV was performed using the QIIME2 naive Bayes classifier, trained with the SILVA 138 SSU Ref NR 99 database (34). The information of ITS and metagenomes, including the host species, the design of experiment, and the data accession were recorded.

For the Lab Test Module, the performances and data of ladybirds under different treatments were recorded, including the development days of each stage, the survival condition of each individual, and the sex and weight of adult. For the life history data, the average and standard error for the development time of each stage and the weight of adults, and the survival rates were calculated in R 4.2.1.

### 2.2. Tool data collection and analysis

For the Morphology Tool, the morphological characters, the distributions, and the diet were available for species detections. For morphological characters, some characters convenient for observation and are often used in classification were adopted, including adult body size, color of head and elytra, and whether with hair overall the body. For adult body size, we recorded the shortest and longest measurements of body length and width reported for each species. For color of head and elytra, all colors with records were included.

For the Barcode Tool, reference barcode databases for species detection (COI, COII, 12S rRNA, 16S rRNA, ND2, and CytB) were established under “Analysis-Species detect.” Two types of databases were created: (1) ALL, containing all sequences downloaded from NCBI prior to March 5, 2025; and (2) Accurate, which excluded problematic sequences (e.g., reversed, low coverage, or contaminated). Mitochondrial genomes from eight ladybird species plus *Tribolium castaneum* and *Dastarcus helophoroides* were used to extract barcodes. Alignments were performed with alignseq.pl in PhyloAln (35) using codon mode (invertebrate mitochondrion) except for 12S and 16S rRNA (standard DNA mode). Alignments were refined with PhyloAln and trimmed with trim_matrix.py to retain sequences overlapping ≥80% with the reference. Genetic distances were calculated using MEGA v6 (36).

For the Gene Tool, ladybird-specific nucleotide and protein databases were constructed from the assembled genomes using makeblastdb.

For the Expression Tool, all transcriptome projects described in the Genomics Module were included and separated into five main treatments based on the designs of the experiments, including stage-specific, tissue-specific, food-stress, bacterial injection stress, and others. The differentially expressed genes were classified using the raw expression of each projects by DESeq2 (37) in R 4.2.1.

For the Primer Tool, published and unpublished primers used in the researches were summarized, including the target species, the target regions, and sequences.

### 2.3. Database implementation

LadybirdBase was implemented on a Nginx web server running Ubuntu, with a MySQL database for data storage. The platform architecture was built using Django, with front-end design in HTML, CSS, and JavaScript. Data visualizations employed ECharts and Chart.js. Sequence alignments for species and gene detection were executed with NCBI BLAST+ (38).

## 3. Database contents and usage

### 3.1. Modules

The Biology Module covers three subfamilies, 36 tribes, and 6,872 species (Figure 1). A phylogenetic tree at the tribal level, based on Huang et al. (24), is displayed (Figure 2A). Users can click on each node to access classification details (e.g., synonyms, Chinese names), morphological traits, diets, distributions, omics data, and references (Figure 2B). Bubble sizes indicate the number of related records, linking to detailed information pages.

**Figure 2.**
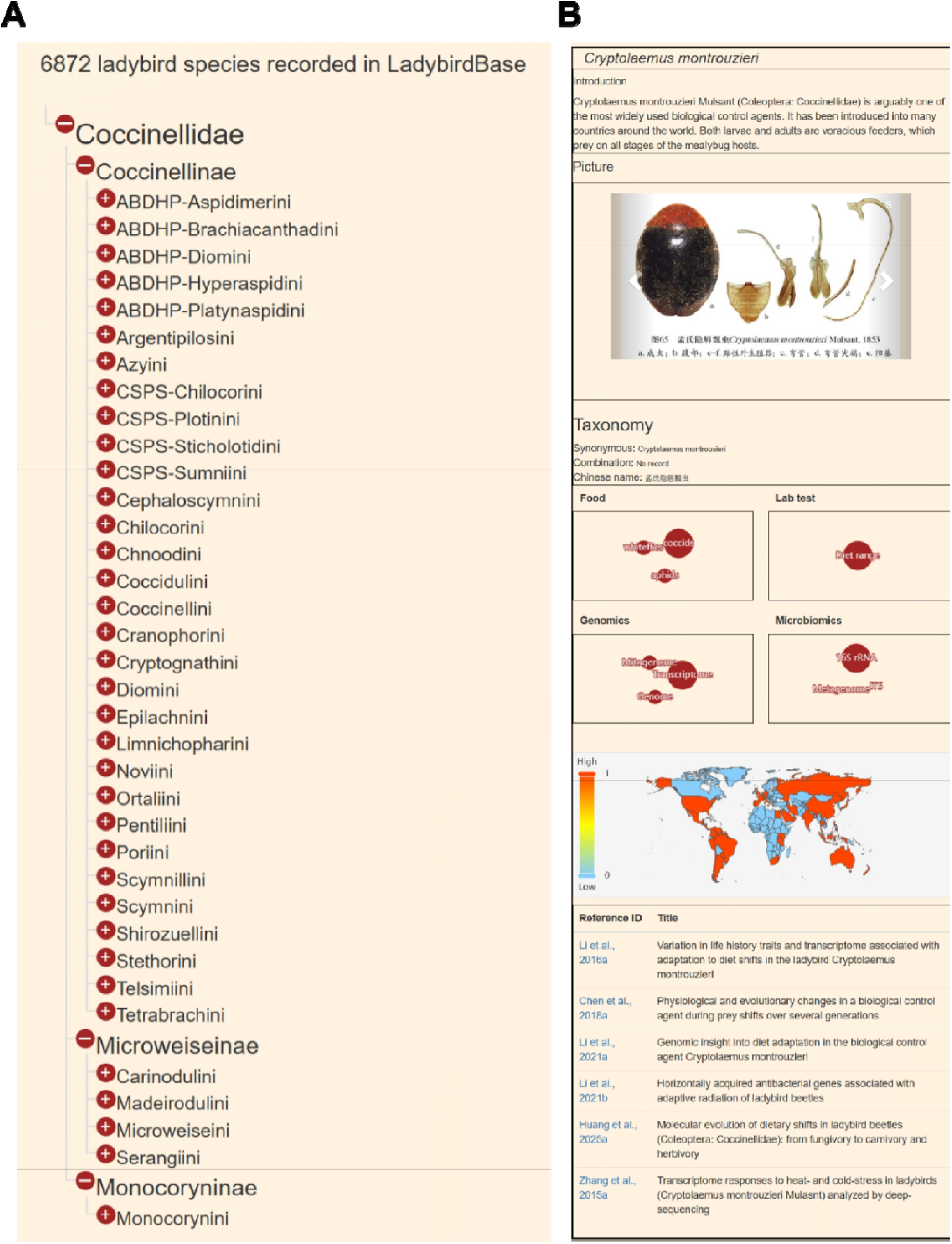
Overview and utilization of the ‘Biology’ module in LadybirdBase. (A) The composition of species information; (B) Species interface.

The Ecology Module is divided into two sections—Diet and Distribution (Figure 3). In Diet Section, ten prey types and five predator species used in biological control are listed (Figure 3A). For each prey type, diet types, relevant ladybird records, feeding distribution, and references are provided. For each predator species, prey types, feeding distribution, and references are available (Figure 3A). In Distribution Section, 10,109 records across 174 countries are displayed (Figure 3B). Country/province-specific details mirror the Biology Module format.

**Figure 3.**
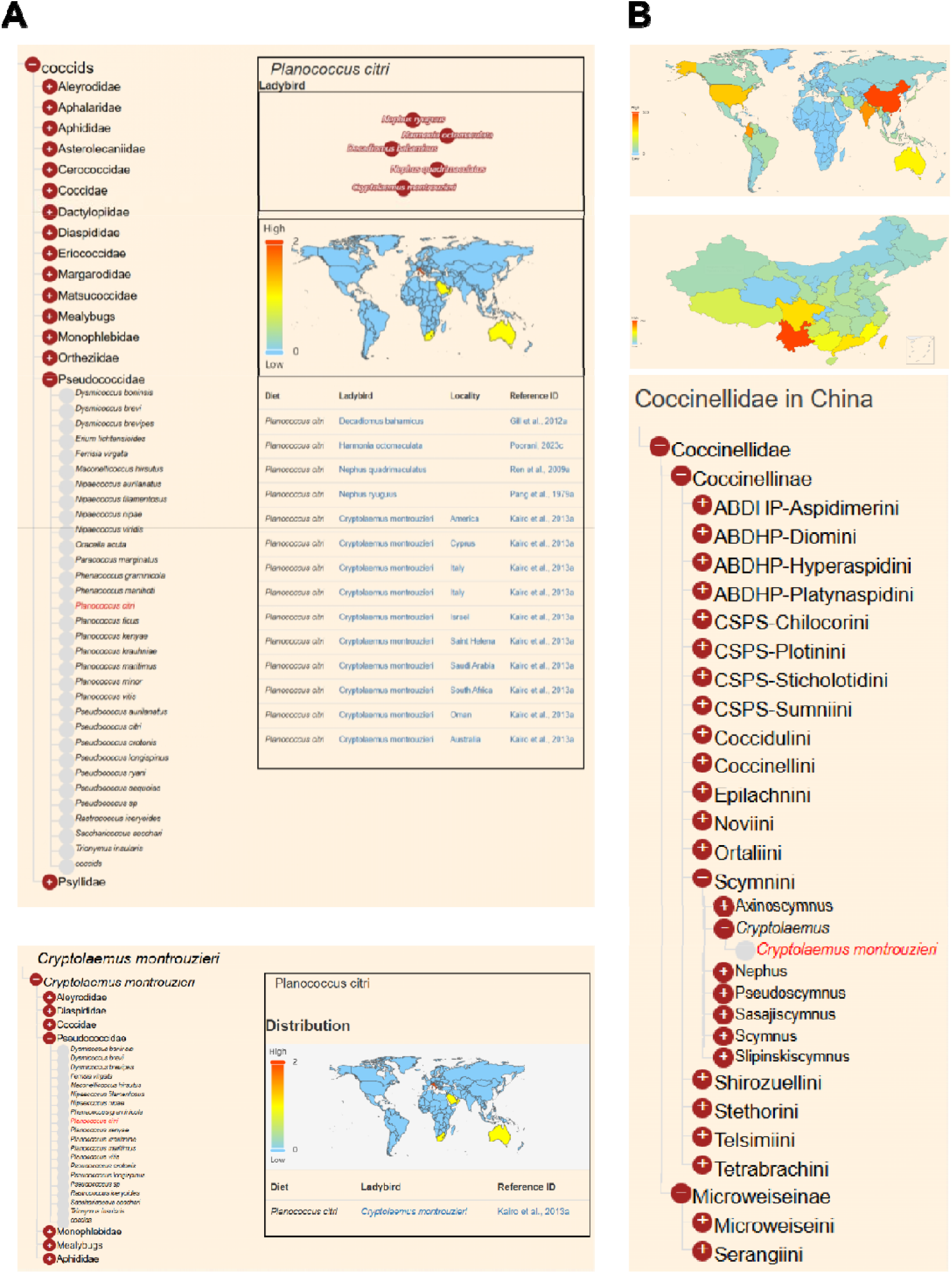
Overview and utilization of the ‘Ecology’ module in LadybirdBase. (A) Diet interface by choosing preys or predators; (B) Distribution interface.

The Genomics Module is divided into five sections. The Whole Genome Section contains 20 assembled genomes (Figure 4A), with details on assembly level, coverage, length, scaffold/contig numbers, N50, GC content, and gene/protein counts. The Transcriptome Section summarizes 782 transcriptomes from 17 species (Figure 4A), including treatments and standardized naming conventions. The Mitogenome Section lists size, annotated genes, accession numbers, and references for 66 species (Figure 4A). The Orthogroup Section features orthogroup analysis from Huang et al. (24), highlighting 32 gene families related to chemosensation, digestion, detoxification, and immunity (Figure 4B), plus 24 horizontally transferred genes (Figure 4B). All orthogroups are annotated using the KEGG KO database, with gene-level details (annotation, expression, variation) accessible (Figure 4B). The Variance Section offers genome-wide variation datasets from two unpublished *C. montrouzieri* resequencing projects: Project 1 (comparison among lab-reared individuals, 21,857,600 SNPs and 4,805,518 InDels) and Project 2 (comparison among wild-collected individuals, 25,231,129 SNPs and 7,639,879 InDels). Variants include position, alleles, consequence type, effect, and genotype distributions.

**Figure 4.**
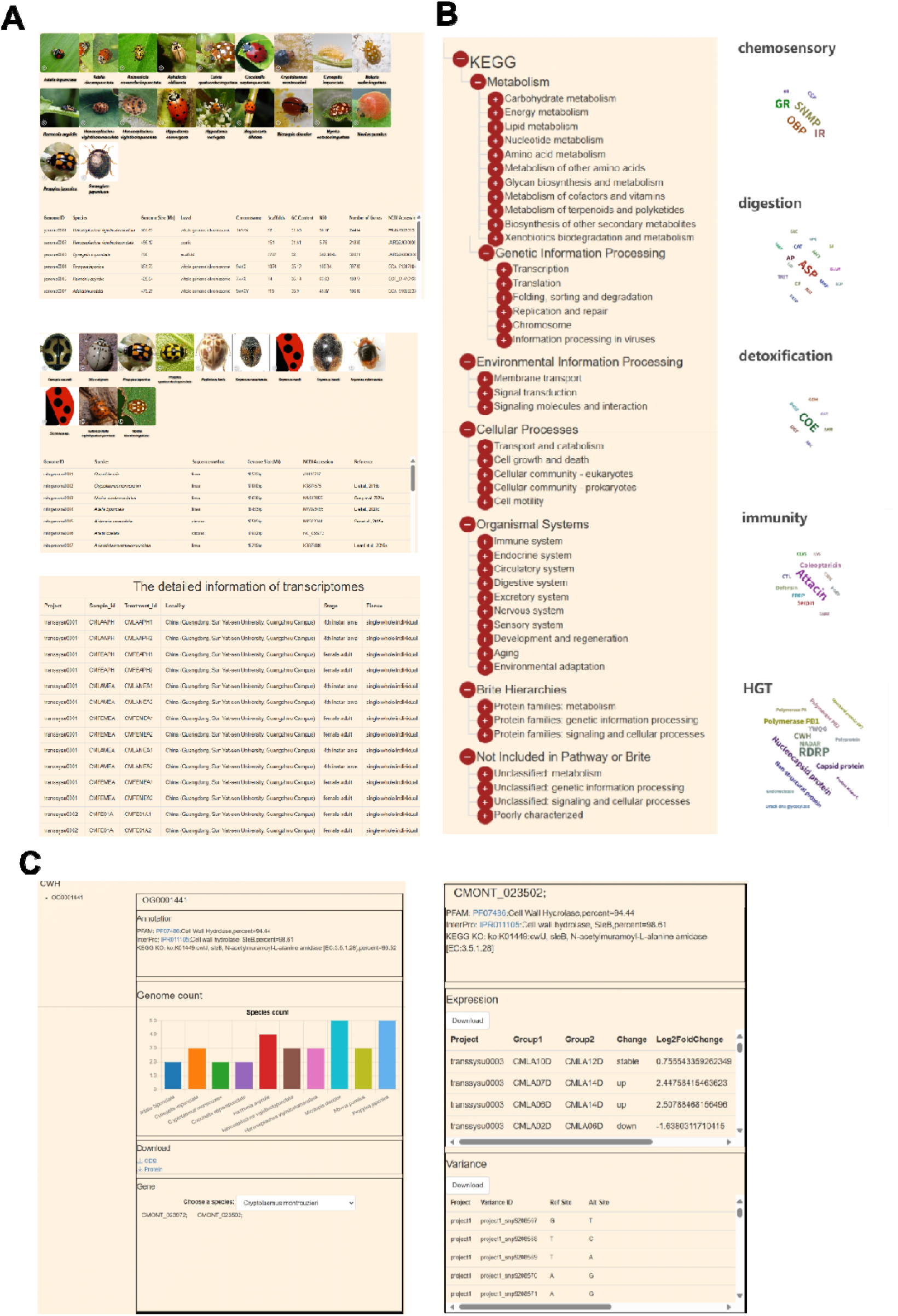
Overview of the ‘Genomics’ module in LadybirdBase. (A) Genome interface, including whole genomes, transcriptomes, and mitogenomes; (B) Orthogroup interface displaying by the annotations of KEGG pathways or detected gene families; (C) The result of choosing single orthogroup.

The Microbiomics Module includes 16S rRNA (970 records from 17 species; Figure 5A), with metadata such as location, treatments, and primer sequences, plus full taxonomic annotation (Figure 5A). The ITS section holds 65 records (Figure 5B), and the Metagenome section contains 10 unpublished datasets (Figure 5C), with species, experiment design, and accession details.

**Figure 5.**
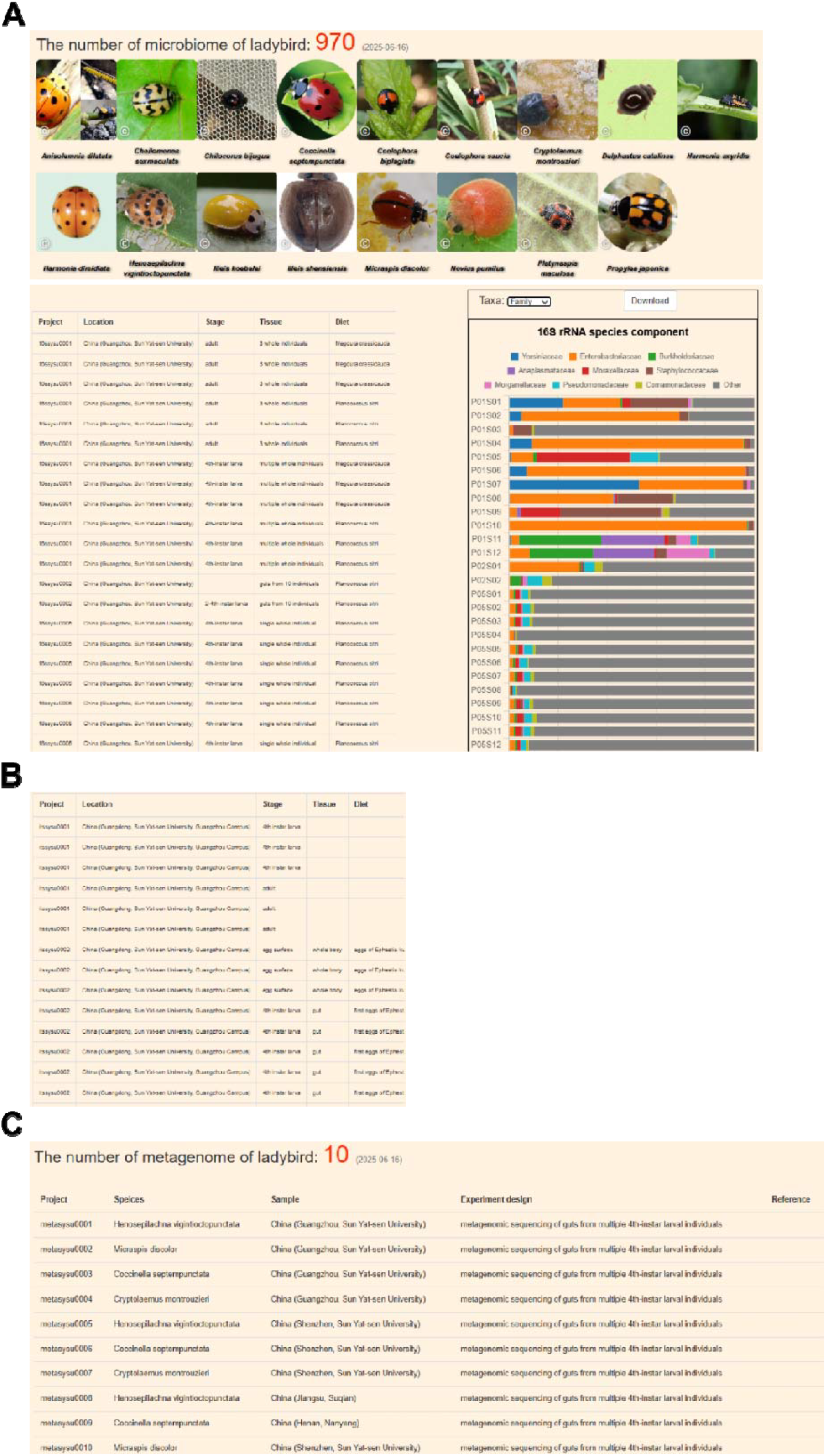
Overview of the ‘Microbiomics’ module in LadybirdBase. (A) 16S rRNA interface. The information of data and the composition of taxonomy are displayed; (B) ITS amplicon interface; (C) Metagenome interface.

The Lab Tests Module is divided into three sections. The Diet Range Section covers lab feeding tests for 154 species and 10 prey types (Figure 6A), showing feeding, development, and reproduction records. The Life History Section includes published and in-house experimental data for 7 key species, with treatments, populations, and results on development time, survival, and adult weight (Figure 6B). The Stress Tolerance Section summarizes pesticide effects on 6 species (41 insecticides), detailing type, concentration, target species, and impact (e.g., survival, development, egg hatch) (Figure 6C).

**Figure 6.**
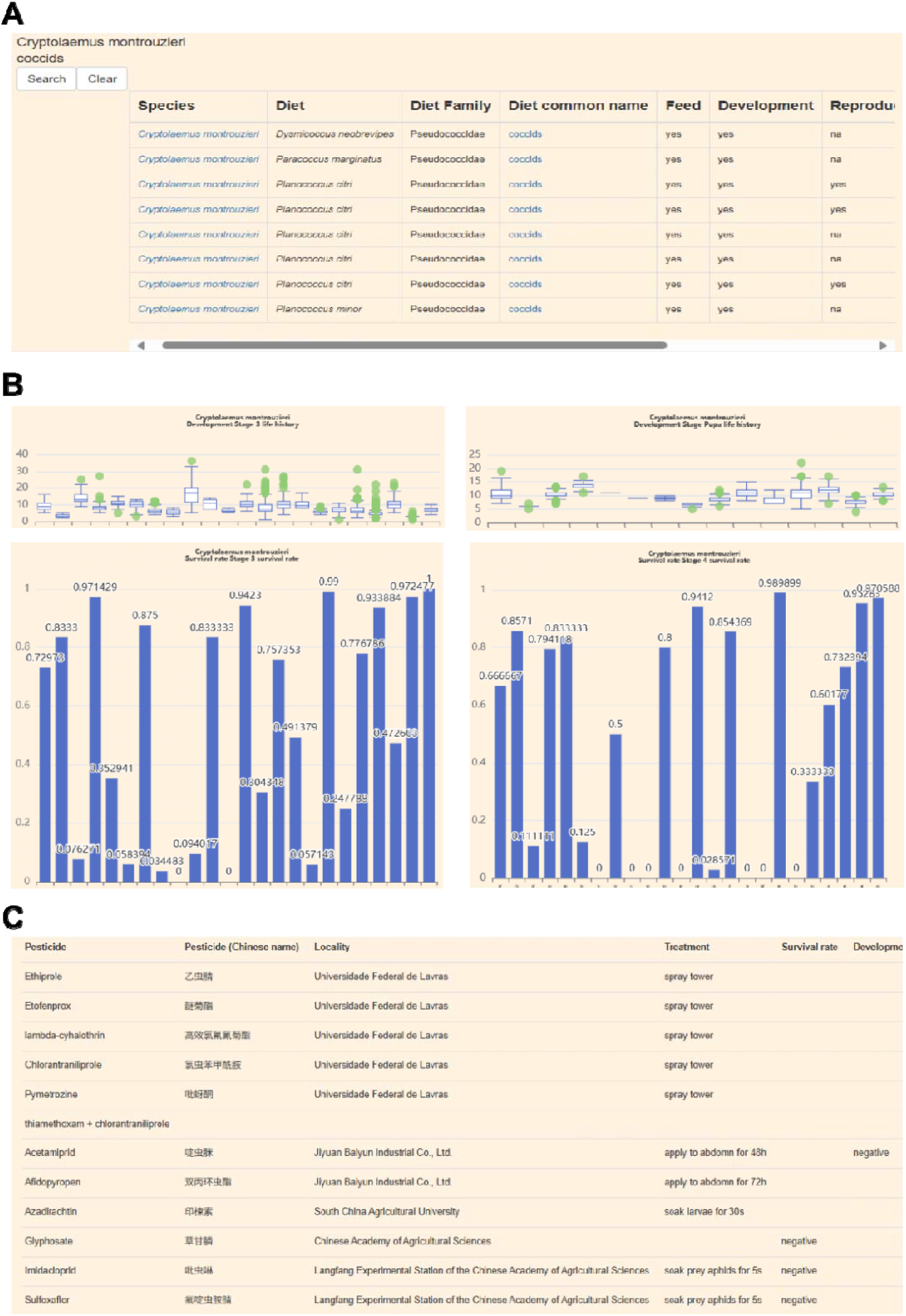
Overview and utilization of the ‘Lab test’ module in LadybirdBase. (A) Diet range interface. The composition of 737 records of food test under the laboratory condition are displayed; (B) Life history interface. The information of life history with different treatments are displayed. (C) Stress tolerance interface. The information of ladybird, including the type of pesticides and the performance of ladybird are displayed.

### 3.2. Tools

The Morphology Tool Identifies probable species by selecting diet, distribution, and morphology, using an in-house database of 864 records (Figure 7A). The Barcode Tool allows barcode sequence alignment against two databases—”All” (12,032 sequences, 835 species) and “Accurate” (9,080 sequences, 614 species)—using six common markers (COI, COII, ND2, CytB, 12S rRNA, 16S rRNA). The Gene Tool aligns gene sequences against LadybirdBase’s genome-based protein/nucleotide databases, with direct links to expression data. The Expression Tool includes differential gene expression analysis from 61 transcriptome projects across 17 species, divided by treatments (Figure 7B). Results include gene/isoform, log2FoldChange, p-value, orthogroup, and functions, with data sortable by change or function. The Primer Tool includes a total of 312 primer pairs targeting 85 regions, which makes it convenient to use directly without the need for repeated design and verification (Figure 7C).

**Figure 7.**
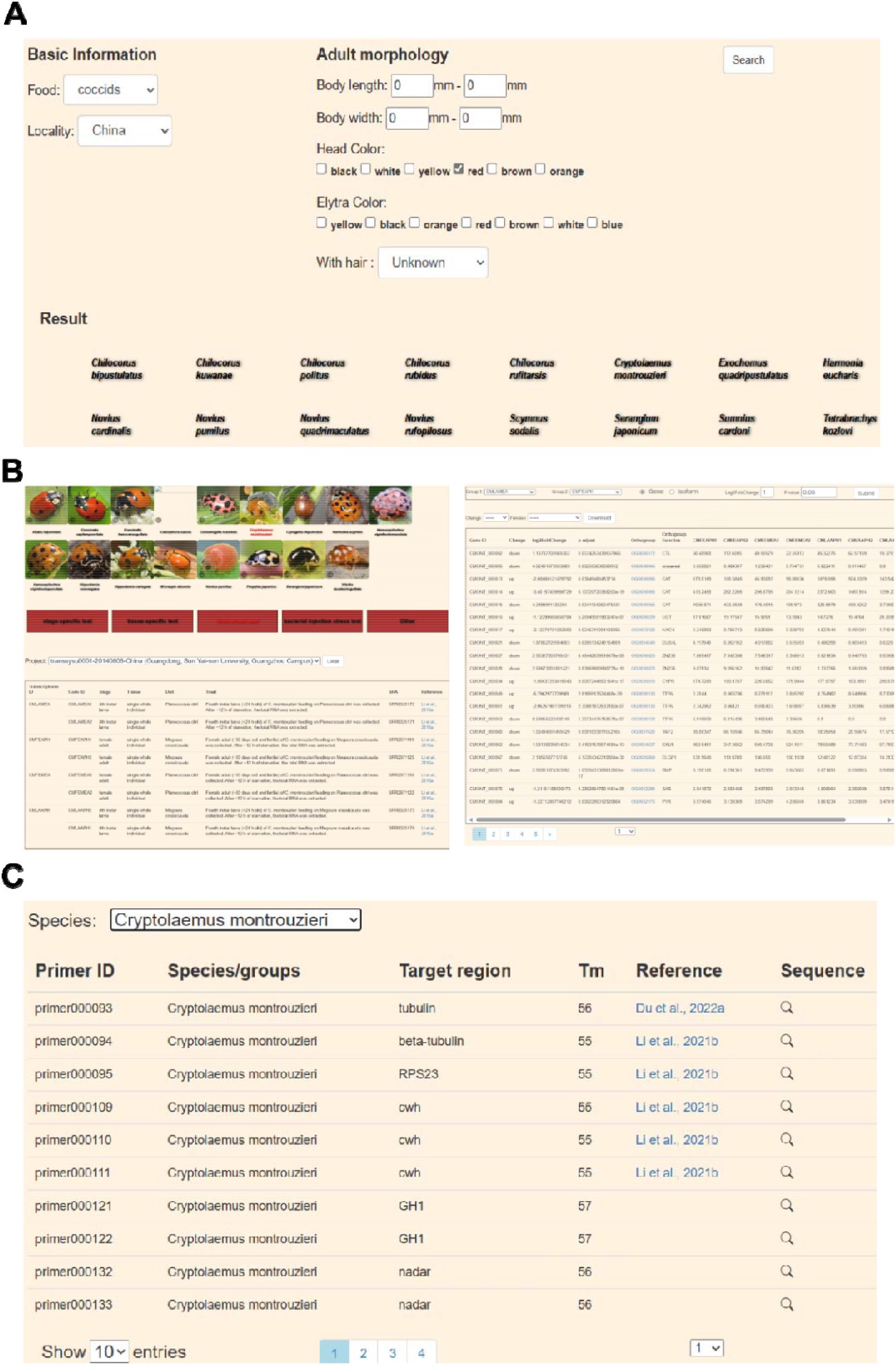
Overview of the ‘Barcode’ Tool in LadybirdBase. (A) The process of species detection with distribution, diet, and morphology information; (B) The process of exploring the gene expression of specific species or treatment; (C) The process of finding specific primers for specific species.

## 4. Discussion and Future Development

Ladybirds are important both in biological control and as model organisms, and the rapid advancement of sequencing technologies has enabled a “next-generation biological control” approach (10). Despite a long history of Coccinellidae use in biocontrol, the lack of a comprehensive, standardized database has hindered fully exploiting their potential. LadybirdBase fills this gap as the first dedicated resource for Coccinellidae by encompassing a variety of biology, ecology and omics data. A key strength of LadybirdBase is its comprehensive omics integration. It provides easy access to raw counts, differential expression, and orthogroup annotations, facilitating exploration of gene functions. Standardized orthologues simplify cross-study comparisons. The microbiomics module, built on a standardized pipeline, ensures accurate and comparable data across treatments. LadybirdBase also manually compiles laboratory test results, including diet ranges essential for selecting effective biocontrol species. Visualized life history data support developing cheaper, alternative feeds. The stress tolerance module helps address non-target effects of pesticides, vital for practical pest management. Finally, LadybirdBase offers user-friendly tools for species detection, sequence comparison, and DGE analysis. Users can leverage ecological data or barcode sequences for species identification, while the expression tool supports quick screening of candidate genes for specific traits.

In the future, LadybirdBase will continue to improve by expanding with new data, including genomic and microbiomic data; adding RNAi targeted sequences; developing tools with features such as species detection from pictures; and constructing AI models capable of generating responses to users’ questions.

## Data availability

LadybirdBase can be accessed at http://www.ladybirdbase.com.

## Author contributions

S.-Y.F., L.-H.S. and P.-H. designed the project. S.-Y.F. and L.-H.S. managed and coordinated the project. S.-Y.F., Y.-K.Y., L.-H., L.-Y.S., C.-L.Q., X.-J.Y., Z.-Y.W., L.-J.Y., M.-Q., W.-Y.M., C.-D., and Q.-M.X. collected datasets. S.-Y.F. and L.-H.S. performed the bioinformatics analysis. S.-Y.F., L.-H.S., and L.-C.R.A wrote the manuscript.

